# Targeting CoV-2 Spike RBD and ACE-2 Interaction with Flavonoids of Anatolian Propolis by *in silico* and *in vitro* Studies in terms of possible COVID-19 therapeutics

**DOI:** 10.1101/2021.02.22.432207

**Authors:** Halil Ibrahim Guler, Fulya Ay Sal, Zehra Can, Yakup Kara, Oktay Yildiz, Ali Osman Belduz, Sabriye Çanakci, Sevgi Kolayli

## Abstract

Propolis is a multi-functional bee product with a rich in polyphenols. In this study, the inhibition effect of Anatolian propolis against SARS coronavirus-2 (SARS CoV-2) was investigated as *in vitro* and *in silico*. Raw and commercial of propolis samples were used in the study and it was found that both of were rich in caffeic acid, p-coumaric acid, ferulic acid, t-cinnamic acid, hesperetin, chrysin, pinocembrin and caffeic acid phenethyl ester (CAPE) by HPLC-UV analysis. The ethanolic propolis extracts (EPE) were used in the screening ELISA test against the spike S1 protein (SARS Cov-2): ACE-2 inhibition KIT for *in vitro* study. Binding energy constants of these polyphenols to the CoV-2 Spike S1 RBD and ACE-2proteinwere calculated separately as molecular docking study using AutoDock 4.2 molecular docking software. In addition, pharmacokinetics and drug-likeness properties of these eight polyphenols were calculated according to the SwissADME tool. Binding energy constant of pinocembrin was the highest for both of the receptors, followed by chrysin, CAPE and hesperetin. *In silico* ADME behavior of the eight polyphenols were found potential ability to work effectively as novel drugs. The findings of both studies showed that propolis has a high inhibitory potential against Covid-19 virus. However, further studies are needed.

## 1. INTRODUCTION

The severe acute respiratory syndrome (SARS) coronavirus-2 (SARS CoV-2) is responsible for coronavirus (COVID-19) pandemic. Coronavirus family has seventh member that infect human beings after SARS coronavirus and Middle East respiratory syndrome (MERS) coronavirus (Bachevski et al., 2020; Zhu et al., 2020). Compared with other viruses, this virus has a high transmissibility and infectivity and is mostly spread by respiratory tracts. The virus is transmitted directly or indirectly, mostly through mucous membranes, nose, mouth and eyes. Until an effective vaccine or medicine is found, many physical and chemical solutions are used to protect against this virus. Mask, distance and hygiene are the most widely used physical protective agents. There are some natural food supplements and vitamins are used in strengthening the immune system. The most used of them are vitamins D, C and propolis (Bachevski et al. 2020; Scorza et al., 2020).

Propolis is a resinous honeybee product obtained from beehives as raw. Honeybees collect propolis mostly from the leaves, barks and trunks of the trees, then transform it with some secretions and store it in the hive. Honeybees benefit from propolis in physical, chemical and biological aspects (Bankova et al., 2019). They used it as antiseptic and antimicrobial, antiviral, antioxidant, antitumoral e.g. agents. Propolis has been also used extensively in traditional and complementary medicine for these wide biological activities (Pasupuleti et al., 2017). In the last 30 years, pharmacological and biochemical studies showed that propolis has wide biological active properties such as antibacterial, antiviral, anti-inflammatory, antitumoral, hepatoprotective, neuroprotective, enhanced immunity in apitherapeutic applications (Pasupuleti et al., 2017; Bankova et al., 2019; Kolayli et al., 2020).

Although its composition and biological active properties depend on the flora of the area where it is collected, propolis consists of approximately 50% resin and balsam, 30% wax, and the rest of essential oils and aromatic compounds (Bankova et al. 2019; Aliyazicioglu et al., 2011; Kızıltas and Erkan, 2020). The active ingredients of propolis, which consists of approximately 300 different organic compounds, are various polyphenols and volatile compounds found in the balsamic part. Although propolis is partially extracted by dissolving in water, glycol, vegetable oils, the most ideal solvent is 60-70% ethanol (Oroian et al., 2020). Today many different commercial propolis extracts are available in different forms such as drops, sprays, pills, pastil, etc. The higher polyphenols or flavonoids contained propolis samples are accepted the higher qualities (Oroion et al., 2020). Polyphenols are the biggest of phytochemical compounds, and polyphenol-rich diets have been associated with many health benefits. Studies are strongly supports that dietary polyphenols is used in the prevention of degenerative diseases, particularly cardiovascular, neurodegenerative and cancers diseases (Tsao, 2010; Pasupuleti et al., 2017).

Propolis is a good antimicrobial and antiviral natural mixture (Przybyłek and Karpiński, 2019). Many studies have been shown that propolis has an antiviral effect againts various DNA and RNA viruses, such as HIV, *Herpes simplex*, HSV-1, HSV-2, *para*-influenza virus, influenza virus type A B, adenovirus, avian reovirus, Newcastle virus disease, bovine rotavirus, pseudo rabies virus etc. (Bachevski et al., 2020; Bankova et al., 2014). The oldest antiviral activity study of propolis against coronaviruses was conducted in 1990. In an *in vitro* study, only antiviral effects of five propolis flavonoids, chrysin, kaempferol, quercetin, acacetin and galangin were investigated. Among them, quercetin exhibited antiviral activity depending on the dose (Debiaggi et al., 1990).

In the COVID-19 pandemic, propolis and some bee products have renewed interest against SARS CoV-2 infection, and some molecular docking studies have confirmed this. *In silico* studies are reported that some of the active ingredients of propolis, especially some flavonoids, have a higher binding potential than the antiviral drugs (Hydroxychloroquine and Remdesivir) used in COVID-19 spike protein and ACE-2 (Mady et al., 2020; Shaldam et al., 2020; Güler and Kara, 2020; Guler et al., 2020). In these studies, it has been shown that the active components of propolis have high binding potential to cellular Angiotensin-converting enzyme-2 (ACE-2) receptors of the S1 spike protein, serine protease TMPRSS2 and PAK1 signaling pathways (Beratta et al., 2020; Scorza et al., 2020). A clinical study was conducted in which propolis tablets were administered in PCR-positive Covid-19 patients (400 and 800 mg) (3×1) for 7 days together with placebo, the results showed that the propolis reduced hospitalization time (Silveira et al., 2021). Propolis also has immunomodulatory, anti-thrombosis activities (Beratta et al., 2020). These activities are also very important in combating the virus. In addition, propolis has been to inhibit the systemic inflammatory response and protect hepatic and neuronal cells in acute septic shock (Korish and Arafa, 2011).

Although propolis is one of the most commonly used natural prophylactic agents during the pandemic, the scientific studies on propolis are insufficient. Therefore, in this study, the inhibition of Anatolian propolis against COVID-19 virus was investigated for the first time in terms of the spike S1 protein (SARS CoV-2): ACE-2 inhibitor screening ELISA test as an *in vitro* study.

## 2. MATERIALS AND METHODS

### 2.1 Chemicals

ELISA KIT of COVID-19 spike protein:ACE-2 assay kit (Cat. No. 79954) was purchased from BPS Bioscience (79954), San Diego, USA gallic acid, protocatechuic acid p-OH benzoic acid, catechin, caffeic acid, syringic acid, epicatechin, p-coumaric acid, ferulic acid, rutin, myricetin, resveratrol, daidzein, luteolin, t-cinnamic acid, hesperedin, chrysin, pinocembrin, caffeic acid phenethyl ester (CAPE), FeSO_4_.7H_2_O, Folin-Ciocalteu’s phenol, diethyl ether, ethyl acetate, acetonitrile were purchased from Sigma-Aldrich (Chemie, Munich, Germany). Daidzein from Cayman Chemical (Michigan, USA) and Ferric tripyridyl triazine (Fe-III-TPTZ), FeCI_3_, CH_3_CO_2_NaAH_2_O, acetonitrile purchased from Merck (Darmstadt, Germany).

### 2.2 Propolis samples

Two different propolis samples were used in this study. Both propolis are samples of Anatolian flora, one was prepared from raw Anatolia propolis, the second was commercial Anatolia propolis. In order to obtain homogeneous Anatolian propolis sample (P1), propolis samples of seven different regions (Van, Rize, Zonguldak, Mugla, Antalya, Diyarbakır and Giresun) were mixed equally. 3 g of the powdered raw propolis was added 30 mL 70%ethanol and shaken on a shaker at a controlled speed for 24 hours (Heidolp Promax 2020, Schwabach, Germany), and ultrasonic (Everest Ultrasonic, Istanbul, Turkey) extraction has been applied for 30 min at 99% power adjustment, then the mixture was filtered through 0.2 μm cellulose filters (Millipore, Bedford, MA, USA). The ethanolic propolis extract of the second samples elected among commercial propolis samples (P2), was supplied by Bee&You (Bee’O^®^) (SBS Scientific Bio Solutions Inc., Istanbul, Turkey). The commercial propolis extract is sold in pharmacies and is widely used for apitherapeutic purposes in Turkey.

### 2.3 Characterization of the propolis samples

#### 2.3.1 Total phenolic compounds (TPC)

Total phenolic content of the both samples were measured with Folin-Ciocalteu’s test using gallic acid (GA) as standard (Singleton et al., 1999). 20 μL six different propolis extracts and standard samples dilutions (from 0.500 mg/mL to 0.015 mg/ml) and 0.2 N 400 μL Folin reagents were mixed and completed to 5.0 ml with distilled water, then vortexed. After 3 min incubation, 400 mL of Na2CO_3_ (10%) was added and incubated at 25°C. The absorbance was measured at 760 nm after 2 h incubation. The total phenolic content was expressed in mg GAE/mL using a standard curve.

#### 2.3.2 Total Flavonoid Content (TFC)

Total flavonoid concentrations of the propolis samples were measured by spectrophotometric method using quercetin standard (Fukumoto and Mazza, 2000). 250 μL of different propolis extracts and standard dilutions (from 0.500 mg/mL to 0,015 mg/ml), 50μL mL of 10% Al(NO_3_)_3_ and 50μL of 1 M NH_4_.CH_3_COO was added and completed 3.0 mL with methanol (99%), vortexed and incubated at 25°C for 40 min. After incubation, the absorbance was then measured against a blank at 415 nm. The total flavonoid concentration was expressed in mg QUE/ml by the curve.

#### 2.3.3 Determination ferric reducing/antioxidant power (FRAP)

The total antioxidant capacities of the samples were determined by using Ferric reducing/antioxidant power assay (FRAP)(Benzie & Strain, 1999). Firstly, working FRAP reagent (Ferric tripyridyl triazine (Fe-III-TPTZ) was prepared. For this, it was freshly obtained by mixing 300 mM pH: 3.6 acetate buffer, 10 mM TPTZ and 20 mM FeCl_3_ solutions in a ratio of (10: 1: 1). Before the samples test, a standard curve is prepared with 1000 μM stock FeSO_4_.7H_2_O solution by serial dilutions. 1.500 ml the FRAP reagent, 50 μL sample and 50 μL methanol were mixed and incubated for 4 min at 37°C, and the absorbance was read at 595 nm against a reagent blank containing distilled water. FRAP value was expressed in μmol FeSO_4_.7H_2_O equivalents/mL.

#### 2.3.4 Determination of phenolic compositions by HPLC-UV

For preparation of the propolis extracts for chromatographic analysis, 10 ml of ethanolic extracts were evaporated and the residue dissolved using 10 ml of purified water of pH 2.The aqueous solution was extracted three times with 5 ml of diethyl ether (15 min, 200 rpm, 25 °C) and three times with ethyl acetate (15 min, 200 rpm, 25 °C). The organic phase, which was collected in a flask after each extraction, was evaporated. The residue was dissolved in 2 mL of methanol and filtered 0.45 μm filters and given to HPLC device for analysis. The phenolic content analysis of the samples was done in triplicate.

Phenolic content analysis of the samples was performed with 280 nm wavelength in RP-HPLC system (Elite LaChrome; Hitachi, Tokyo, Japan) with C18 column (150 mm * 4.6 mm, 5 μm; Fortis). In the analysis using 70% acetonitrile/water (A) and 2% acetic acid/water (B) as mobile phase, the injection volume was 20 μl, the flow rate was 1.00 ml/min and the column temperature was 30°C. The analysis was done using a gradient program. The R2 values of the calibration curves of the nineteen standard phenolic compounds used in the analysis were between 0.998 and 1.000.

### 2.4. Inhibition Assay for Covid-19

The Spike S1 (SARS CoV-2): ACE-2 inhibitor scanning colorimetric assay kit (Cat. No. 79954) was purchased from BPS Bioscience (79954), San Diego, USA. The colorimetric test is designed for screening and profiling inhibitors of this interaction. The aim of the test is to prevent the virus from being entering the cell by preventing the interaction between Spike protein S1 and ACE-2. Using the kit protocol, the absorbance was read at 450 nm using UV/Vis spectrophotometer microplate reader. The propolis and standard phenolic samples were diluted with 70% ethanol and the Covid-19/ELISA test procedure was applied. All tests were done in triplicate.

### 2.5. Molecular Docking Studies

AutoDock 4.2 software for performing molecular docking studies was used to investigate the possible interactions of eight ligands and reference molecule with the target proteins. To evaluate the prediction of accuracy of binding affinity between ligands and two target proteins, the binding free energies (ΔG) are calculated for the crystal structures and the docking mod. The 3-D structure of all ligands (pinocembrin, chrysin, cape, hesperetin, ferulic acid, t-cinnamic acid, p-coumaric acid, caffeic acid) and reference molecule (Hydroxychloroquine) were retrieved from the PubChem database (https://pubchem.ncbi.nlm.nih.gov/) as sdf format and then converted to pdb format by using BIOVIA DS Visualizer software (Dassault Systèmes BIOVIA, 2016).

### 2.6. Pharmacokinetics and drug-likeness properties (ADME Prediction)

In order for a drug to be effective, it must reach its target in the body in sufficient concentration and remain in bioactive form long enough for the expected biological events to occur there. Drug development involves absorption, distribution, metabolism and excretion (ADME) increasingly earlier stage in the discovery process, at a stage where the compounds are abundant but access to physical samples is limited (Daina et al., 2017). Pharmacokinetics, drug-likeness and medicinal chemistry properties of eight ligands were predicted using the Swiss ADME server. Important parameters related to ADME properties, such as Lipinski’s five rules, drug solubility, pharmacokinetic properties, molar refraction and drug likeliness were analyzed. The SMILES format retrieved from PubChem Database of the interested ligands were used as input for analysis tool (Daina et al., 2017).

### 2.7 Statistical analyses

The statistical evaluations were carried out with the SPSS Statistic 11.5 (IBM SPSS Statistics, Armonk, New York, USA). For presenting the results, descriptive statistics were used as mean ± SD. The correlation analyses were performed with Mann–Whitney U-test. The significance was determined at p < 0.05.

## 3. RESULTS

### 3.1. Propolis analyses

Table 1 shows analysis of the two Anatolian propolis samples. The ethanolic propolis extracts, one was prepared from raw Anatolia propolis (P1) and the other was commercially available (P2). The pH values of both propolis samples were between 4.50 and 4.80, both of the found acidic. The total phenolic substance was found to be 12.30 mg GAE/mL in P1 sample and 40.68 mg GAE/ml in P2 sample. It was found that commercial sample (P2) has 3 times higher phenolic compound (P1). Similar to the total amount of phenolic content, the amount of total flavonoid substance was found to be different in both samples, and it was found to be 12.40 mg / mL in the commercial sample and 1.04 mg / mL in the P1 sample. Total antioxidant capacities of the samples were investigated only FRAP assay, and the results is showed that the commercial sample (P2) was higher two times than the other (P1). The phenolic profile results of two ethanolic propolis samples are summarized in Table 2. In the phenolic composition analyzes performed by HPLC-UV, it was found that both propolis samples contained similar phenolic components, but the flavonoids in the commercial sample (P2) were higher.

**Table 1.** Analysis of two Anatolian propolis samples

|  |  | **pH** | **Total Phenolic**<br>**Content**<br>**(mgGAE/mL)** | **Total Flavanoid**<br>**Content**<br>**(mgQUE/mL)** | **Total Antioxidant**<br>**Capacity (FRAP)**<br>**$\mu\text{molFeSO}_4/\text{mL}$** |
| --- | --- | --- | --- | --- | --- |
| P1 | Raw Propolis<br>(Prepared by self) | 4.80±0.01 | 12.30±0.02 | 1.08±0.03 | 141.40±1.70 |
| P2 | Commercial Propolis<br>(BEE'O)© | 4.50±0.01 | 40.68±1.50 | 12.40±0.98 | 285.40±5.40 |

**Table 2.** Phenolic profile of the EPE samples by HPLC-UV

| Phenolic Standards<br>(mg/ 100 g) | Raw Propolis<br>(P1)<br>(Prepared by self) | Commercial<br>Propolis<br>(BEE'O)© (P2) |
| --- | --- | --- |
| Gallic acid | - | - |
| Protocatechuic acid | - | - |
| *p*-OH Benzoic acid | - | - |
| Catechin | - | - |
| Caffeic acid | 707.71 | 497.27 |
| Syringic acid | - | - |
| Epicatechin | - | - |
| *p*-Coumaric acid | 881.89 | 269.02 |
| Ferulic Acid | 378.53 | 145.77 |
| Rutin | - | - |
| Myricetin | - | - |
| Resveratrol | - | - |
| Daidzein | - | - |
| Luteolin | - | - |
| *t*-Cinnamic acid | 510.47 | 51.75 |
| Hesperetin | 711.06 | 3029.46 |
| Chrysin | 665.11 | 1190.89 |
| Pinocembrin | 1685.48 | 1804.22 |
| CAPE | 3268.72 | 3168.26 |

(-):not detected

### 3.2. Molecular Docking Studies

The structures of the polyphenols used in the molecular docking program are given in Figure 1. The binding free energies values for ACE-2 and SARS-CoV-2 Spike RBD were calculated using the AutoDock 4.2 software program are summarized in Table 3. The docked poses, interacting residues and interactions of each ligand with ACE-2 and SARS CoV-2 Spike RBD are given in Figures 2–9. Details about estimated binding affinities (kcal/mol) and Ki values of docked ligands are shown in the Table 3. The result of the study showed that four (pinocembrin, chrysin, caffeic acid phenethyl ester and hesperetin) were showed very low binding free energies to the ACE-2 receptor and SARS-CoV-2 Spike Protein RBD. It was also found that these four flavonoids have higher binding potential than hydroxychloroquine, which was used as a covid-19 drug and was used as the standard ligand in the study. From the Table 3, it can be clearly predicted that pinocembrin has the highest binding energy value of −8.58 kcal/mol for ACE-2 protein and −7.54 kcal/mol for SARS CoV-2 Spike RBD, followed by chrysin owing dock scores of −8.47 and −7.48 kcal/mol, respectively.

**Fig. 1.**
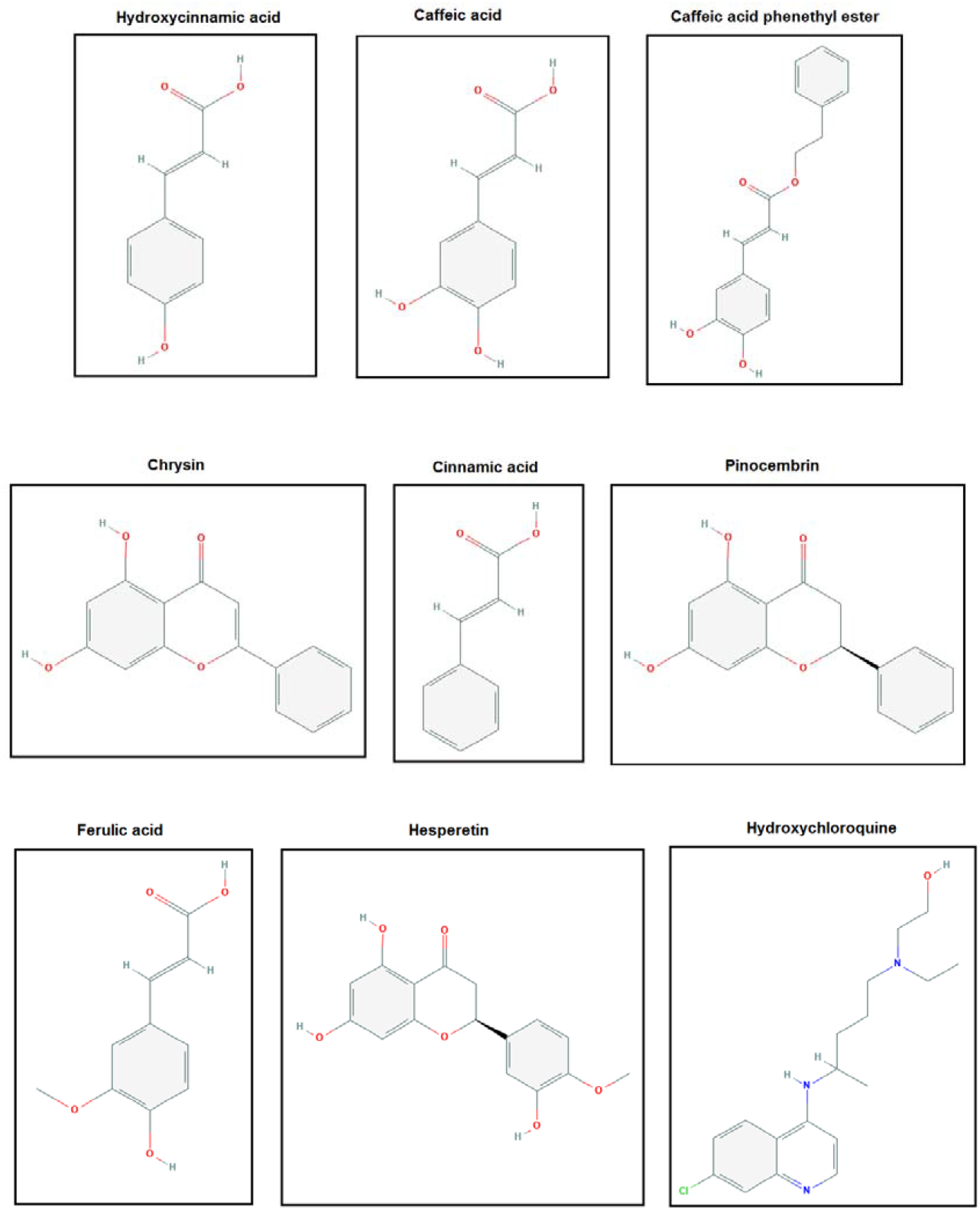
2-D structures of ligands used in the present study

**Fig. 2.**
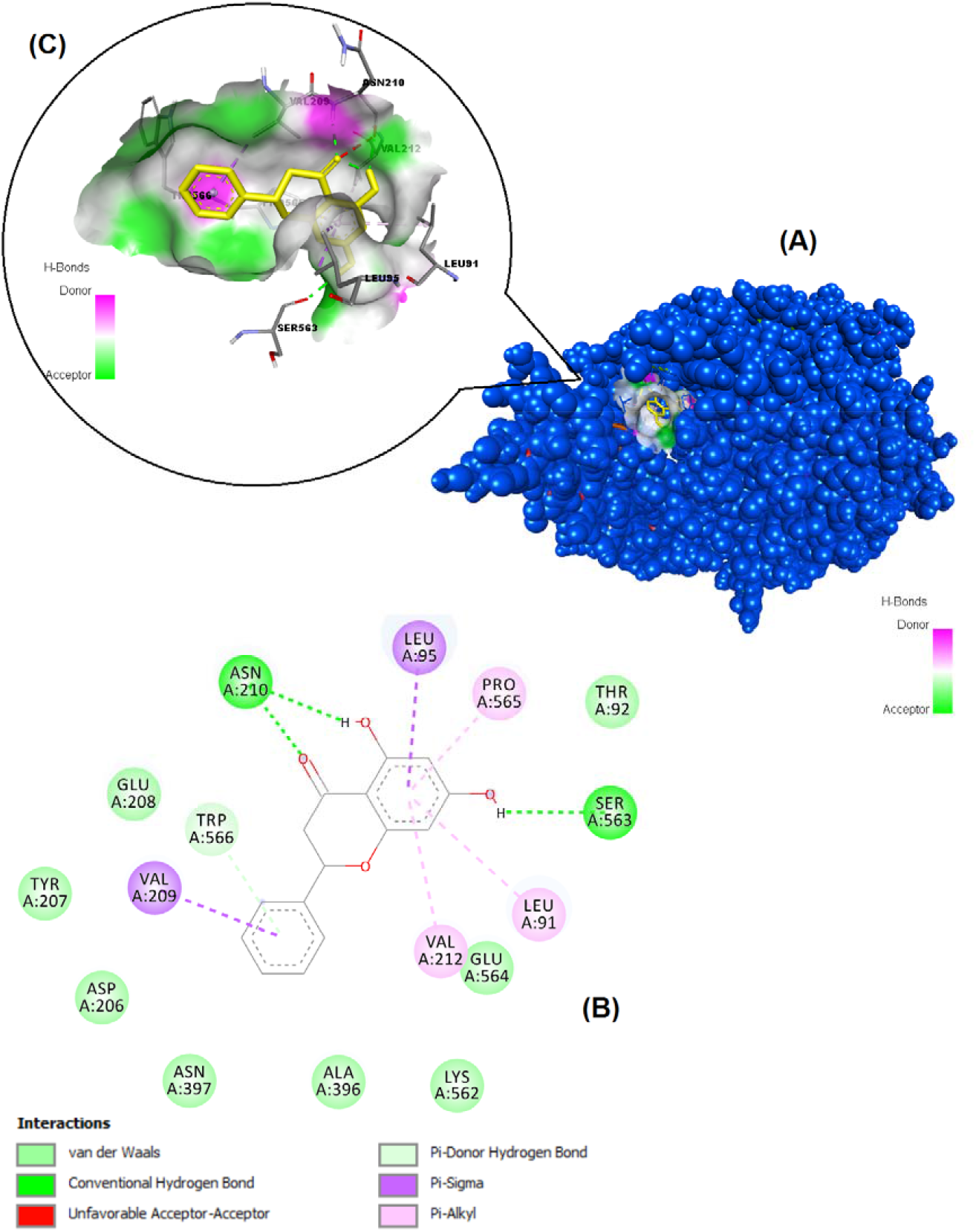
Binding pose profile of pinocembrin in the target protein ACE2 (A), blue shaped molecule represents the receptor and yellow shaped molecule indicates the ligand. The two-dimension (2D) (B) and three-dimension (3D) (C) interactions analysis of ACE2 protein with compound pinocembrin.

**Fig. 3.**
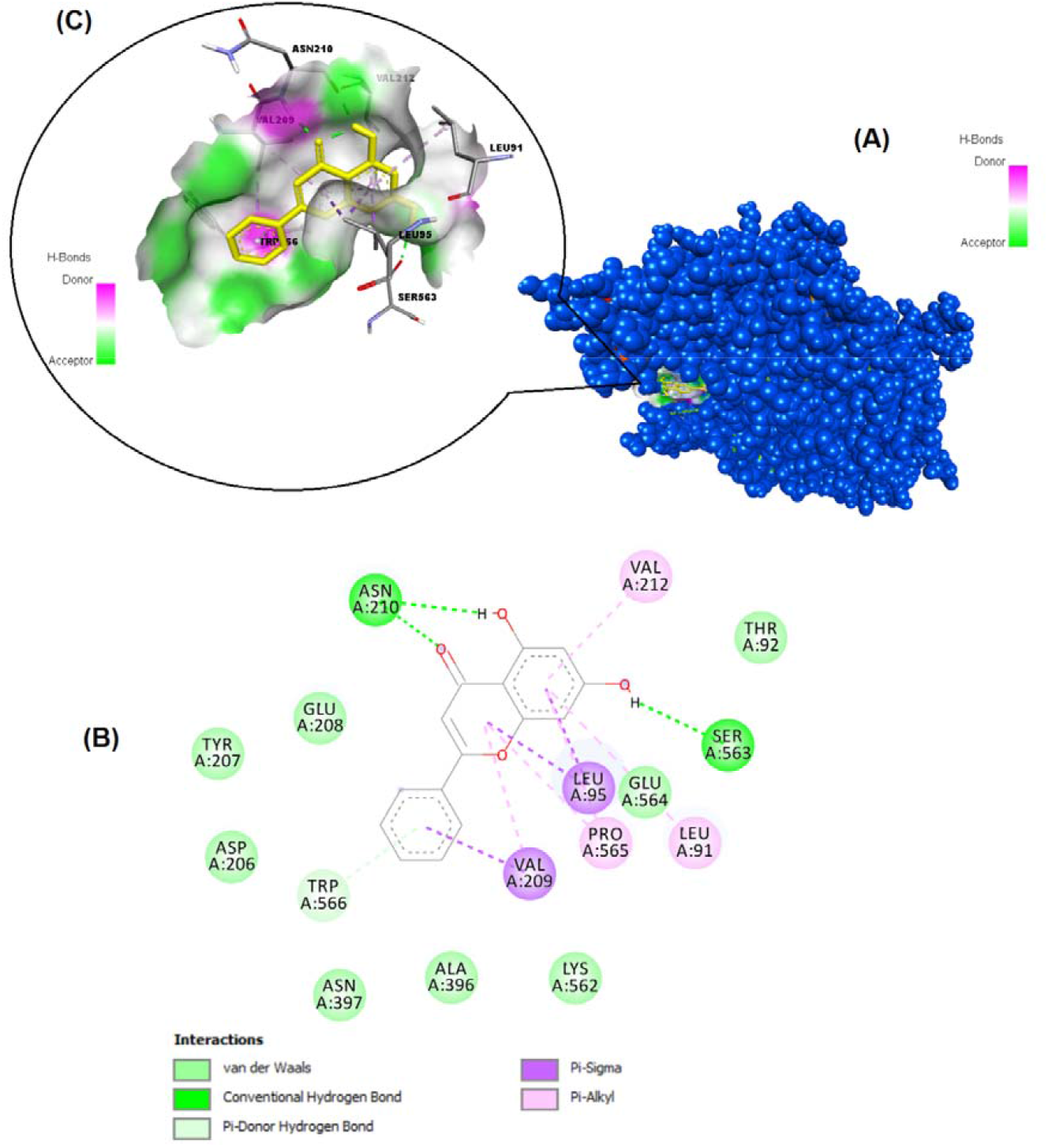
Binding pose profile of chrysin in the target protein ACE2 (A), blue shaped molecule represents the receptor and yellow shaped molecule indicates the ligand. The two-dimension (2D) (B) and three-dimension (3D) (C) interactions analysis of ACE2 protein with compound chrysin.

**Fig. 4.**
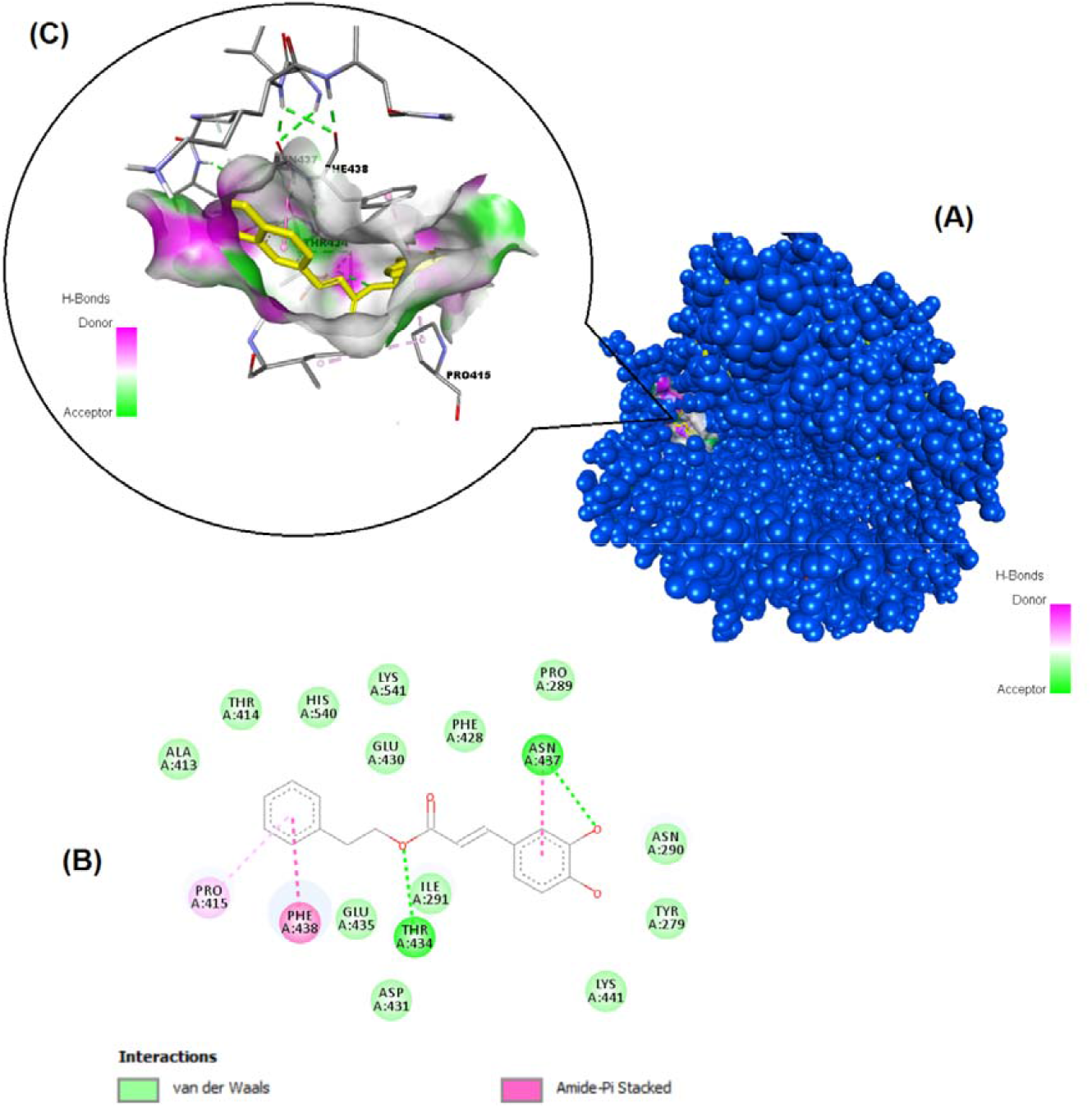
Binding pose profile of CAPE in the target protein ACE2 (A), blue shaped molecule represents the receptor and yellow shaped molecule indicates the ligand. The two-dimension (2D) (B) and three

**Fig. 5.**
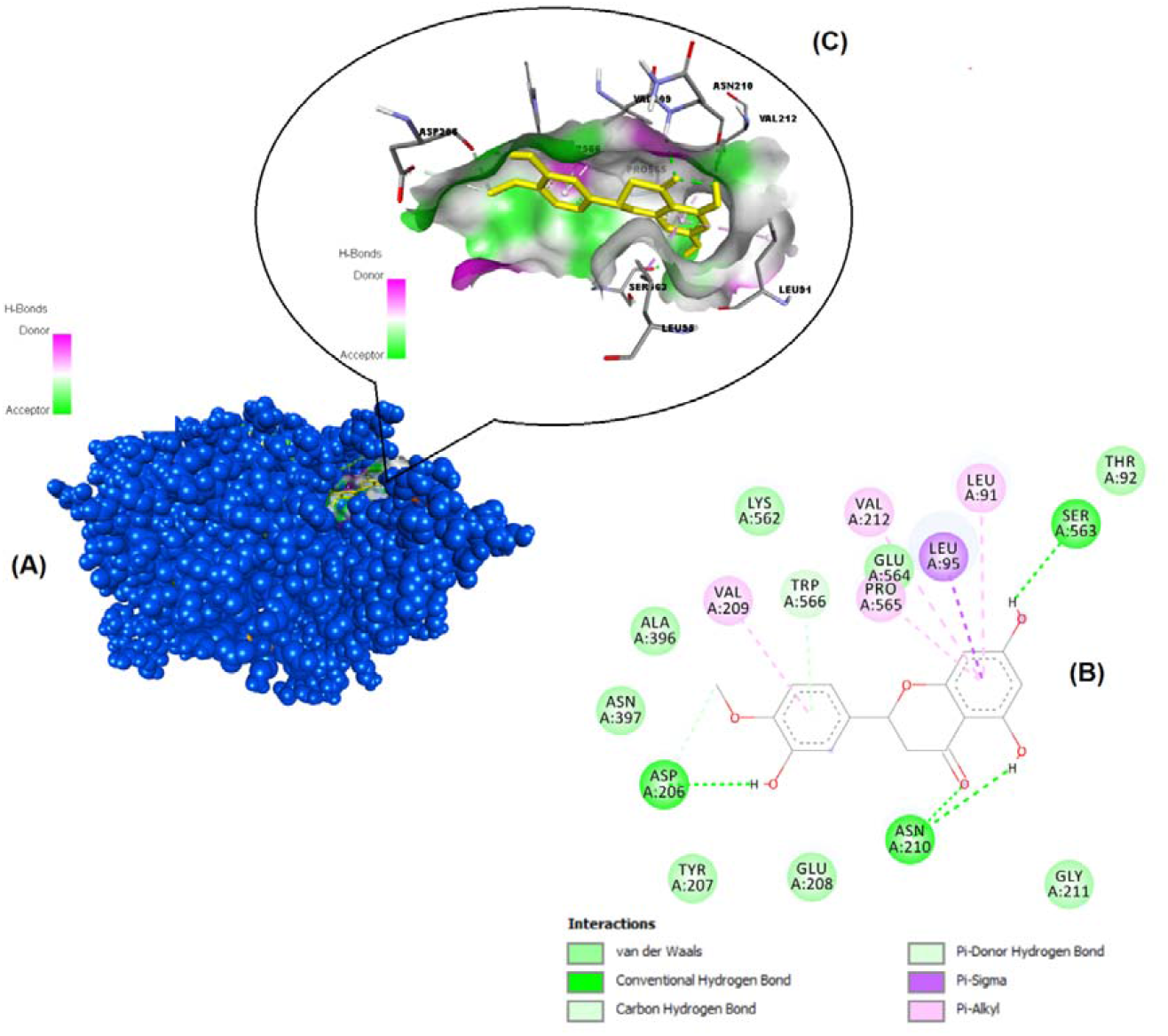
Binding pose profile of hesperetin in the target protein ACE2 (A), blue shaped molecule represents the receptor and yellow shaped molecule indicates the ligand. The two-dimension (2D) (B) and three-dimension (3D) (C) interactions analysis of ACE2 protein with compound hesperetin.

**Fig. 6.**
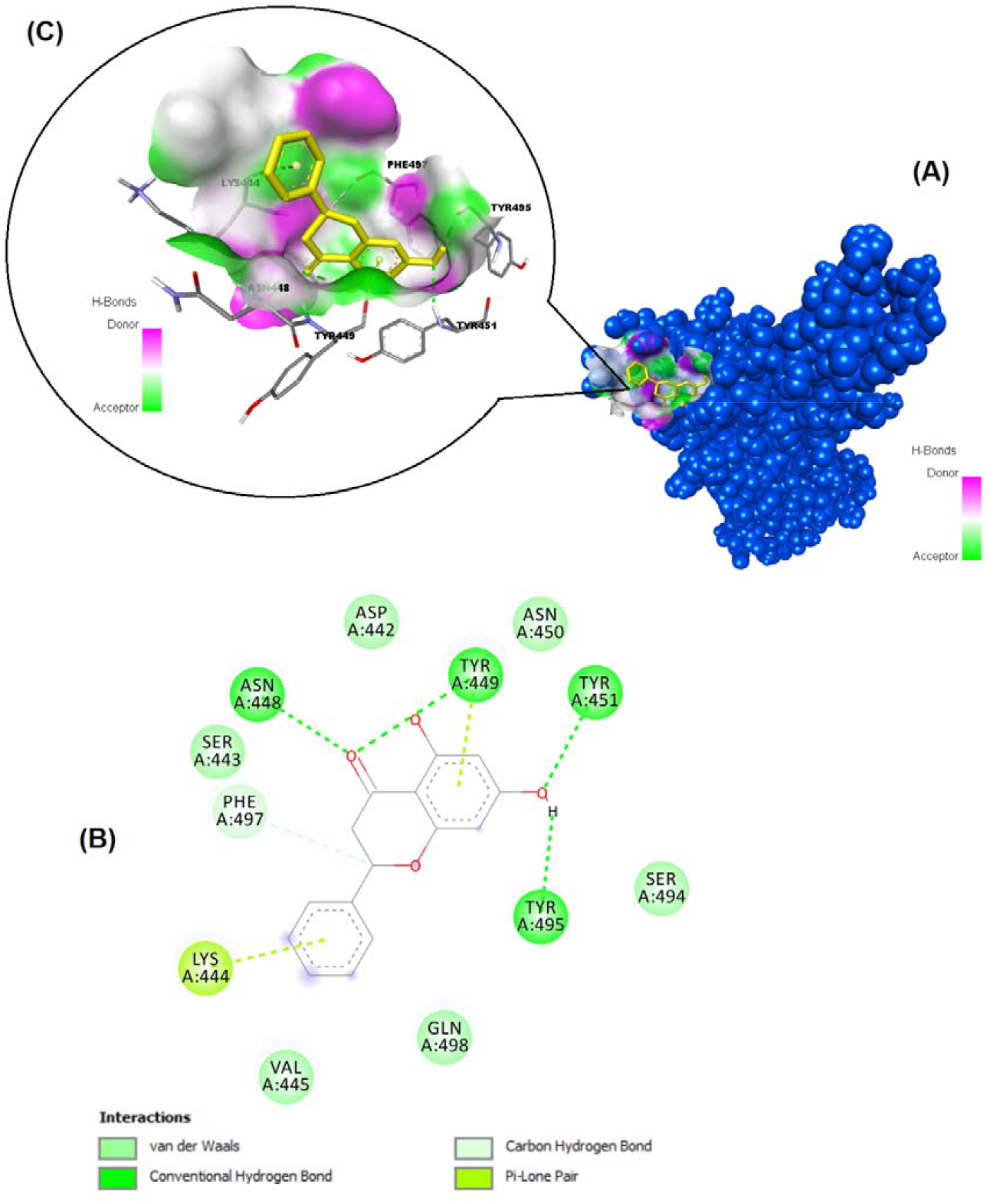
Binding pose profile of pinocembrin in the SARS-CoV-2 Spike receptor binding domain (A), blue shaped molecule represents the receptor and yellow shaped molecule indicates the ligand. The two-dimension (2D) (B) and three-dimension (3D) (C) interactions analysis of SARS-CoV-2 Spike RBD with compound pinocembrin.

**Fig. 7.**
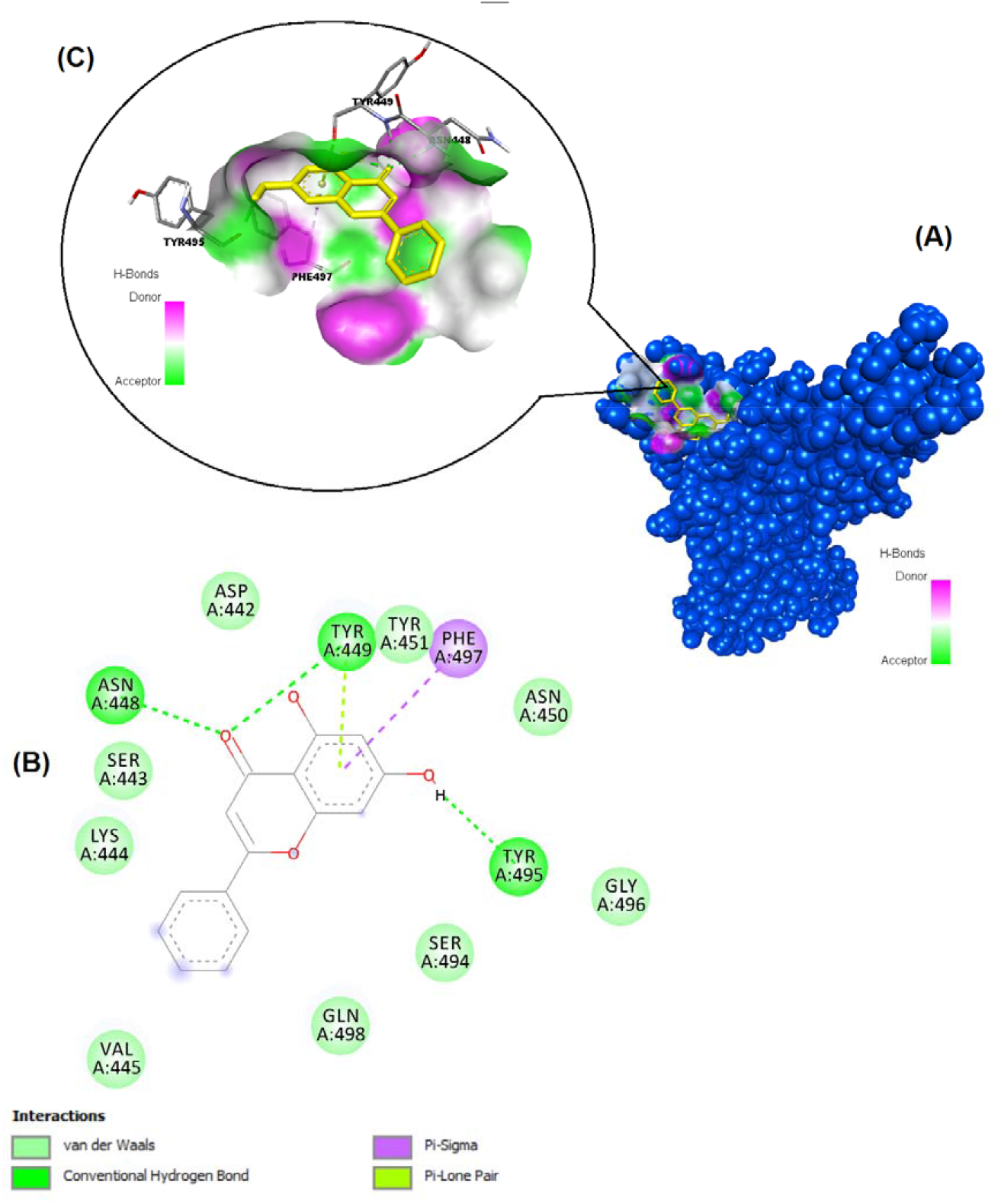
Binding pose profile of chrysin in the SARS-CoV-2 Spike receptor binding domain (A), blue shaped molecule represents the receptor and yellow shaped molecule indicates the ligand. The two-dimension (2D) (B) and three-dimension (3D) (C) interactions analysis of SARS-CoV-2 Spike RBD with compound chrysin.

**Fig. 8.**
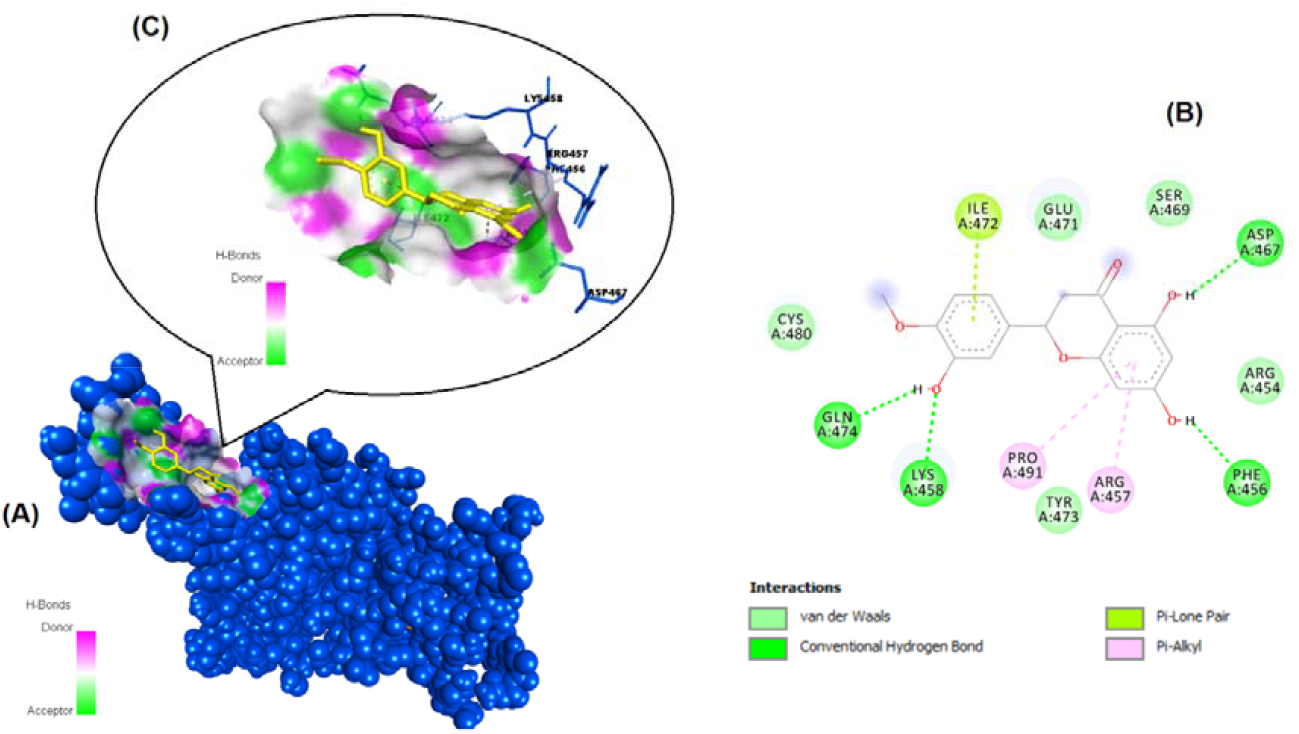
Binding pose profile of hesperetin in the SARS-CoV-2 Spike receptor binding domain (A), blue shaped molecule represents the receptor and yellow shaped molecule indicates the ligand. The two-dimension (2D) (B) and three-dimension (3D) (C) interactions analysis of SARS-CoV-2 Spike RBD with compound hesperetin.

**Fig. 9.**
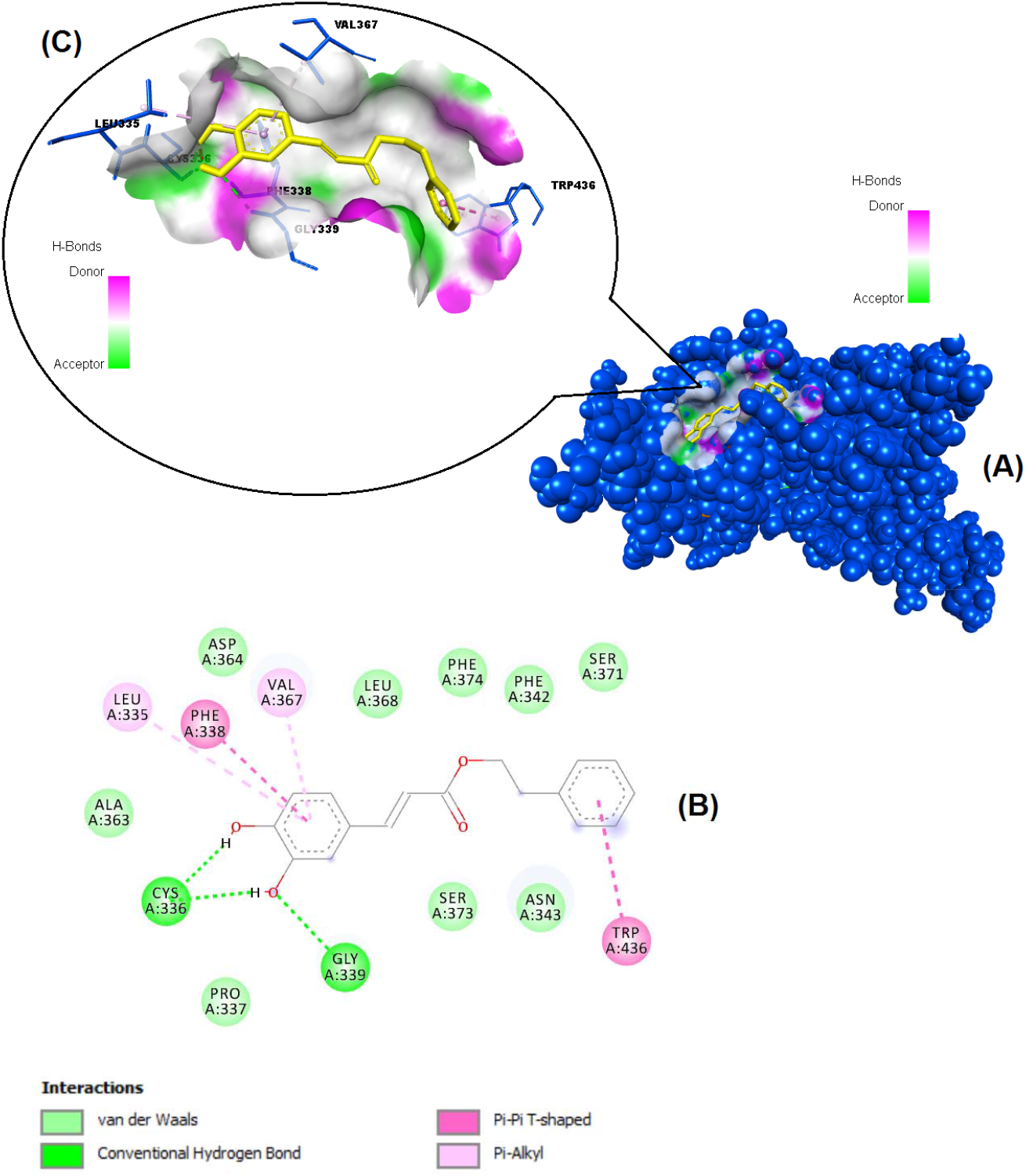
Binding pose profile of CAPE in the SARS-CoV-2 Spike receptor binding domain (A), blue shaped molecule represents the receptor and yellow shaped molecule indicates the ligand. The two-dimension (2D) (B) and three-dimension (3D) (C) interactions analysis of SARS-CoV-2 Spike RBD with compound CAPE.

**Table 3.** Summary of estimated binding affinity (kcal/mol) and Ki values of docked ligands against ACE2 and SARS-CoV-2 Spike receptor binding domain, and interacted residues in the binding sites.

| Receptor Name / PDB ID | Ligand Name | Binding Energy (kcal/mol) | $K_i$ | Interacted residues with ligand |
| --- | --- | --- | --- | --- |
| Angiotensin-Converting Enzyme-2 (ACE-2)<br>EC: 3.4.17.23<br>/<br>6M0J (Chain A)<br>Res: 2.45 Å | Pinocembrin | -8.58 | 510.99 nM | Asn210, Leu9, Pro565, Ser563, Leu91, Val212, Val209 |
|  | Chrysin | -8.47 | 623.53 nM | Asn210, Val212, Ser563, Glu564, Leu91, Leu95, Pro565, Val209 |
|  | CAPE (Caffeic acid phenethyl ester) | -8.42 | 677.67 nM | Asn437, Ile291, Thr434, Phe438, Pro415 |
|  | Hesperetin | -8.22 | 943.94 nM | Leu91, Ser63, Asn210, Asp206, Val209, Trp566, Val212, Glu564, Pro565, Leu95 |
|  | Ferulic acid | -5.65 | 72.03 µM | His540, Ile291, Pro289, Thr434, Glu430 |
|  | t-Cinnamic acid | -5.65 | 72.06 µM | Leu456, Trp477, Leu503, Trp165, Trp271, Lys481 |
|  | p-coumaric acid | -5.63 | 74.21 µM | Trp165, Pro500, Leu503, Leu456, Trp477, Lys481, Trp271 |
|  | Caffeic acid | -5.31 | 127.93 µM | Leu73, Ala99, Leu100, Lys74, Asn103 |
|  | *Hydroxychloroquine | -7.90 | 1.61 µM | Arg393, Phe390, Leu391, Asn394, His378, His401, Asp350 |
| SARS-CoV-2 Spike receptor binding domain<br>/<br>6YLA (Chain A)<br>Res: 2.42 Å | Pinocembrin | -7.54 | 2.99 µM | Asn448, Tyr449, Tyr451, Tyr495, Lys444, Phe497 |
|  | Chrysin | -7.48 | 3.29 µM | Asn448, Tyr449, Phe497, Tyr495 |
|  | Hesperetin | -7.28 | 4.63 µM | Ile472, Asp467, Phe456, Arg457, Pro491, Lys458, Gln474 |
|  | CAPE | -7.17 | 5.54 µM | Leu335, Phe338, Val367, Trp436, Gly339, Cys336 |
|  | Ferulic acid | -6.93 | 8.29 µM | Leu441, Tyr495 |
|  | t-Cinnamic acid | -6.64 | 13.57 µM | Phe497, Lys444, Asn448, Tyr449, Tyr495 |
|  | Caffeic acid | -6.43 | 19.36 µM | Leu441, Tyr495, Phe497 |
|  | p-coumaric acid | -5.97 | 42.06 µM | Phe497, Tyr495, Leu441 |
|  | *Hydroxychloroquine | -6.32 | 23.35 µM | Leu517, Tyr396, Val382, Phe392, Thr430, Phe515 |

\*reference molecule

### 3.3. Pharmacokinetics and drug-likeness properties (ADME Prediction)

Table 4 shows the ADME properties of the polyphenols detected in the propolis samples. According to Lipinski, a compound to be used should have 5 properties to be selected as a potential drug. These are: (a) Molecular mass <500 Daltons (b) high lipophilicity (expressed as LogP 5) (c) less than 5 hydrogen bond donors (d) less than 10 hydrogen bond acceptors (e) molar refractivity between 40 and 130. The scanned eight flavonoid compounds used in this study were all found to meet the Lipinski’s five rules (Table 4). And also, other properties like pharmaco-kinetic, physicochemical and drug-likeness properties are given in Table 4. The results were indicated that all of eight molecules have the potential to work effectively as novel drugs.

**Table 4.** ADME properties of ligands docked with SARS-CoV-2 Spike RBD and ACE-2 target proteins

| Ligand name | (Lipinski's Rule of Five) |  |  |  |  | Heavy atoms | Aromatic heavy atoms | Rotat. bonds | TPSA | ESOL Class | GI absorption | BBB permeant | Pgp substrate | Bio Avail. Score | PAINS alerts | Synthetic Accessibility | Violations | Drug Likeliness |
| --- | --- | --- | --- | --- | --- | --- | --- | --- | --- | --- | --- | --- | --- | --- | --- | --- | --- | --- |
|  | Mol. weight | LogP | H-bond donor | H-bond acceptor | Molar Refractivity |  |  |  |  |  |  |  |  |  |  |  |  |  |
| Pinocembrin | 256.25 | 2.26 | 2 | 4 | 69.55 | 19 | 12 | 1 | 66.76 Å | soluble | High | Yes | No | 0.55 | 0 | 2.96 | No | Yes |
| Chrysin | 254.24 | 2.5 | 2 | 4 | 71.97 | 19 | 16 | 1 | 70.67 Å | moderately soluble | High | Yes | No | 0.55 | 0 | 2.93 | No | Yes |
| CAPE | 284.31 | 3.26 | 2 | 4 | 80.77 | 21 | 12 | 6 | 66.76 Å | moderately soluble | High | Yes | No | 0.55 | 1 | 2.64 | No | Yes |
| Hesperetin | 302.28 | 1.91 | 3 | 6 | 78.06 | 22 | 12 | 2 | 66.76 Å | soluble | High | No | Yes | 0.55 | 0 | 3.22 | No | Yes |
| Ferulic acid | 194.18 | 1.36 | 2 | 4 | 51.63 | 14 | 6 | 3 | 66.76 Å | soluble | High | Yes | No | 0.85 | 0 | 1.93 | No | Yes |
| t-Cinnamic acid | 148.16 | 1.79 | 1 | 2 | 43.11 | 11 | 6 | 2 | 37.30 Å | soluble | High | Yes | No | 0.85 | 0 | 1.67 | No | Yes |
| p-coumaric acid | 164.16 | 1.26 | 2 | 3 | 45.13 | 12 | 6 | 2 | 57.53 Å | soluble | High | Yes | No | 0.85 | 0 | 1.61 | No | Yes |
| Caffeic acid | 180.16 | 0.93 | 3 | 4 | 47.16 | 13 | 6 | 2 | 77.76 Å | very soluble | High | Yes | No | 0.56 | 1 | 1.81 | No | Yes |

**Lipinski's Rule of Five:** Molecular weight (<500 Da), LogP (<5) , H-bond donor (<5), H-bond acceptor (<10), Molar Refractivity (40-130)

### 3.4. *In vitro* inhibition studies

The binding of ACE-2 protein to SARS CoV-2 Spike S1 protein was studied for both EPEs with the inhibitor screening colorimetric assay kit (BPS Bioscience, 79954). The key to this ELISA assay is the high sensitivity of detection of ACE-2-Biotin protein by Streptavidin-HRP. This technique is based on the binding of the active ingredients of the propolis to this spike S1 protein/ACE-2 complex and preventing the binding of the enzyme-labeled second antibody to the protein. The presence of enzyme activity (horseradish peroxidase) indicates the absence of binding. The inhibition values are expressed in terms of the IC_50_ value, and are expressed as the amount of the propolis that provides 50% inhibition. The data are shown in Figures 10–11. Together with the propolis samples, ability of certain flavonoids to inhibit the interaction of SARS CoV-2 S1 spike protein and ACE-2 were also tested. It was found that the two EPE samples were found to cause inhibition of interaction of SARS CoV-2 S1 spike protein: ACE-2 receptors and the degree of inhibition (IC_50_) varied depending on the propolis concentration. The IC_50_ value of the commercial propolis was found to be about 3 times higher than P1 sample.

**Fig. 10.**
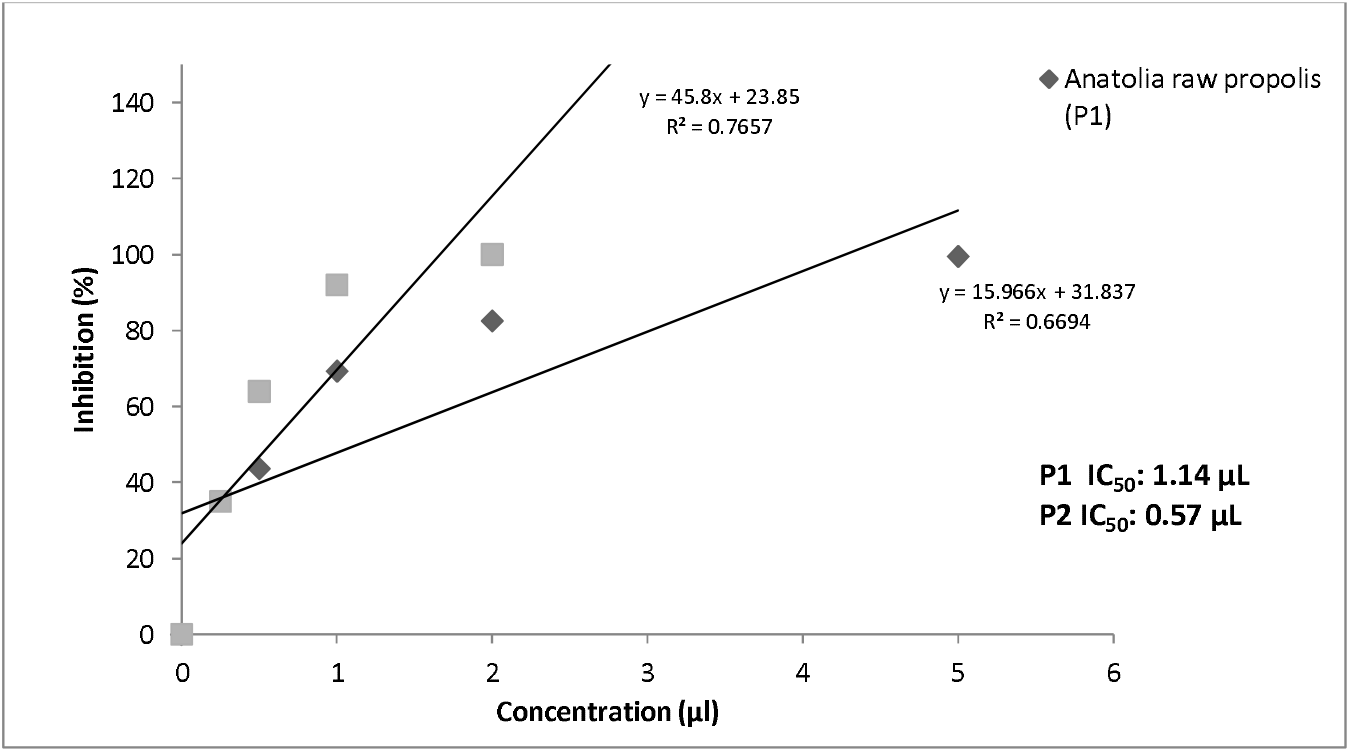
IC_50_ values of P1 and P2 samples

**Fig. 11.**
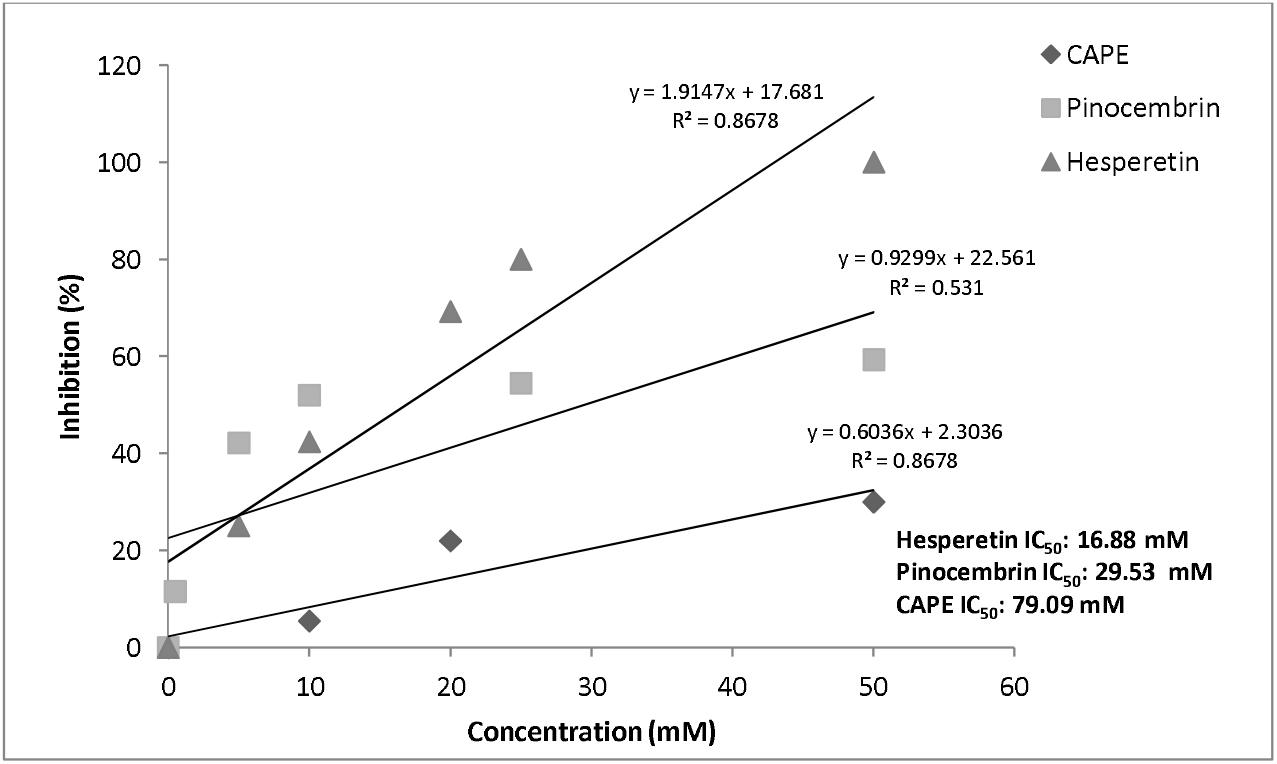
IC_50_ values of CAPE, pinocembrin and hesperetin

In the study, inhibition effect of five different concentrations of hesperetin, CAPE and pinocembrin were tested with the ELISA plate assay was. It was determined that the inhibition values were varied depending on the concentration (Fig 10). Hesperetin is the best inhibitor againts the SARS CoV-2 S1 spike protein and ACE-2, has the lowest IC_50_ value (16.88 mM), pinocembrin and CAPE were followed them. When comparing in silico and *in vitro* study results, it is found that pinocembrin had a high inhibitory effect in *in silico* study, whereas hesperetin was more active in *in vitro* study.

## 4. DISCUSSION

Propolis is a natural bee product and a very good source of polyphenols. The ideal extraction solvent for propolis rich in phenolic acid and flavonoids is 70% ethanol. The most characteristic analysis parameters for propolis are total polyphenol, total flavonoid and phenolic content analysis. It was determined that the phenolic amount of the two ethanolic propolis samples were found similar in terms of their composition, but different in terms of amounts. The main reason for this difference is the amount of the raw propolis that used initially when extracting with solvent (70%). In the P1 sample, 3 g of raw propolis was prepared at a ratio of 1:10 in 30 mL 70% alcohol. However, since the P2 sample is commercial, it is not known how much raw propolis is used; it is only possible to say that it is used in higher amounts than P1 sample. In short, the extract prepared with a high amount of raw propolis is expected to have a higher amount of phenolic matter. However, the quality of raw propolis used in extraction is also important parameter (Yeo et al., 2015). It has been reported that total polyphenol content (TPC) in Anatolian raw propolis samples varied from 115 mg GAE/g to 210 mg GAE/g (Aliyazicioglu et al., 2011). It was reported that TPC is varied from 55.75 to 91.32 mg GAE/g in Brazilian propolis (Andrade et al., 2017) and TPC was varied from 10 to 80 mg GAE/g in Azerbaijan propolis (Zehra et al., 2015). These results show that the amount of total polyphenol in propolis quality is a critical criterion and at the same time, this quality depends on the flora.

The samples were also found acidic, (pH <6.0) that the acidity are sourced from organic acid that were found in the propolis. Caffeic acid, p-coumaric acid, and ferulic acids were detected both of the samples. However, it was reported that gallic acid, caffeic acid, coumaric acid, ferulic acid and syringic acid, protocathequic acid are major phenolic acids in propolis samples (Yeo et al., 2015; Aliyazicioglu et al. 2011). Gallic acid, protocathequic acid, p-OH benzoic acid, syringic acid were not detected in any sample, but this does not mean that there will be no gallic acid in Anatolian propolis (Keskin et al., 2019). Because these phenolic acids are highly polar compounds, they may not have switched to ethanol with a lower polarity than water. CAPE, pinocembrin and chrysin were found the most abundant flavonoids in both of the samples. CAPE is a polyphenol found mostly in propolis, and its high amount reflects the quality of propolis and possessed a wide biological active properties such as antioxidant, anti-inflammatory, anti-tumoral etc. (Aliyazicioglu et al., 2011; Bankova et al., 2014; Bankova et al., 2019; Venkateswara et al., 2017).

Pinocembrin, hesperetin and chrysin were also found abundant flavonoids in the EPEs. Flavonoids are the most common and the largest plant polyphenolic obtained from the everyday plant-source diet (Chun et al., 2007), and have been proven to be responsible for a variety of biological activities such as antioxidant, antibacterial, antiviral and antiinflammatory activity. Amount of flavonoids taken in the daily diet was estimated to be about 200 mg/day, and it consisted of 84% flavan-3-ols, flavanones (7.6%), flavonols (7%), anthocyanidins (1.6%), flavones (0.8%) and isoflavones (0.6%). However, epidemiological studies conducted in populations fed with flavonoid-rich diets have shown that the incidence of cardiovascular damage and some types of it has decreased (Cui et al., 2008). It has been supported by studies that propolis is a very good source of flavonoids (Venkateswara et al., 2017; Kowacz and Pollack, 2020). Thus, high polyphenols and flavonoids are taken together with consumption of propolis as a food supplement.

The antioxidant capacity of the EPEs was measured according to the FRAP test, and this test is a very simple and easy test that shows the total antioxidant capacity. The higher the FRAP value in the analysis measured according to the reduction ability of the Fe (III) TPTZ complex, the higher the antioxidant capacity (Can et al., 2015). It was determined that the antioxidant capacity of the commercial propolis sample (P2) was approximately 2 times higher than the other sample (P1). The high antioxidant capacity in the commercial propolis sample is thought to be due to the high polyphenol content (Can et al. 2015; Kolayli et al., 2020).

Molecular docking is a crucial tool for exploring the interactions between the target protein and a small molecule. The binding energy (kcal/mol) data allows us to study and compare the binding affinity of different ligands/compounds with their corresponding target receptor molecule. The lower binding energy indicates a higher affinity of the ligand for the receptor. The ligand with the highest affinity can be selected as a potential drug for further studies. For this study, eight flavonoids with a broad range of biological activities, along with hydroxychloroquine which exhibited efficacy against SARS CoV-2, have been selected as ligands to investigate their binding affinities with SARS CoV-2 Spike Protein RBD and ACE-2 as target receptor proteins. All the eight polyphenols and one reference molecule were individually docked to the ACE-2 and SARS CoV-2 Spike RBD, respectively. After successful docking of all the ligands used in these docking experiments, the results showed us significant interactions of the ligands with the target receptors. Four ligands (pinocembrin, chrysin, CAPE and hesperetin) are bound to the target protein ACE-2 more effectively than the reference molecule. And also, seven ligands (pinocembrin, chrysin, hesperetin, CAPE, ferulic acid, t-cinnamic acid and caffeic acid) are bound stronger to the SARS CoV-2 spike RBD than the reference molecule, hydroxychloroquine. The docking study results that pinocembrin has the highest binding energy value of −8.58 kcal/mol for ACE-2 protein and −7.54 kcal/mol for SARS CoV-2 Spike RBD, followed by chrysin owing dock scores of −8.47 and −7.48 kcal/mol, respectively. In molecular docking studies with propolis and Covid-19, it has been reported that some propolis flavonoids have high binding affinities to ACE-2 receptors and the virus spike protein. For example, Quercetin and rutin have been reported to have similar activity (Basu et al., 2020; Guler et al., 2020; Guler and Kara, 2020; Mady et al., 2020; Berretta et al., 2020).

The rule essentially determines the molecular properties of a compound that are its primary requirement for being a potential drug, such as absorption, distribution, metabolism, and excretion (ADME). Generally, some parameters are used to evaluate potential interactions between drug and other non-drug target molecules (Das, et al., 2020; Gupta et al.,2020; Jayaram et al., 2012; Lipinski, 2004). The propensity for a compound with a certain pharmacological or biological activity to be used as a potential drug is evaluated. It was found that these eight polyphenols detected in these propolis samples were meet the five rules of Lipinski and other features were also compactable. Hence, we suggest that the flavonoids have the potential to work effectively as novel drugs.

The aim of the study was to investigate the inhibition potential of ethanolic propolis extracts by binding to SARS CoV-2 Spike protein and ACE-2, so two propolis extracts were used in the study. In *in vitro* study, it was found that both samples caused inhibition, but the P2 sample showed higher activity. This is thought to be due to the higher polyphenol content of the P2 sample. The result that the 3 different polyphenols (hesperetin, pinocembrin and CAPE) studied separately cause inhibition of the virus shows us that the active substances in propolis are mostly due to flavonoids. Since no in vitro study was reported before, we could not compare and discuss our results. All eight phenolic standards were not tested as *in vitro* by the ELISA KIT assay, since the plate was limited to 96 well plates. As a result of *in silico* study, the *in vitro* inhibitions of three flavonoids with the highest binding potential were examined.

When comparing *in silico* study results with *in vitro* study results, it is seen that pinocembrin, hesperetin and CAPE have high binding affinities to virus spike S1 protein and ACE-2 receptor. So, *in silico* and *in vitro* studies support each other. In addition, the fact is that the phenolic standards used in terms of ADME properties were found to have high drug properties and the results proved that propolis has high potential in combating the Covid-19.

## 5. CONCLUSION

In this study, it was shown for the first time that ethanolic Anatolia propolis extracts inhibit Covid-19 virus in terms of binding spike S1 protein and ACE-2 receptor as both *in vitro* and *in silico* studies. However, more detailed studies are needed.

## Acknowledgments

This work was supported by the KTU BAP FBA-2020-9192 project. We thank (Bee&You (Bee’O^®^) (SBS Scientific Bio Solutions Inc., Istanbul, Turkey) company for allowing the use of commercial propolis sample.

## Notes

**Conflict of interest**, No conflict of interest is declared.

### Competing Interest Statement

The authors have declared no competing interest.

### Summary of Updates

Results and and discussion section was seperated and revised.

